# Extracellular electron transfer in cable bacteria enables growth rates comparable to aerobic respiration

**DOI:** 10.1101/2025.10.08.681138

**Authors:** Kartik Aiyer, Leonid Digel, Markéta Linhartová, Andreas Schramm, Ugo Marzocchi, Lars Peter Nielsen, Ian PG Marshall

**Affiliations:** Center for Electromicrobiology, Department of Biology, Aarhus University, Aarhus, Denmark; Yusuf Hamied Department of Chemistry, University of Cambridge, Cambridge, U.K.; Center for Water Technology (WATEC), Aarhus University, Aarhus, Denmark

**Author notes:** Corresponding authors: Kartik Aiyer and Ian PG Marshall.

**Keywords:** cable bacteria, extracellular electron transfer, electrode, voltammetry, metalloprotein, bioelectrochemical system

## Abstract

Cable bacteria are filamentous sulphide-oxidisers performing long-distance electron transfer in redox-stratified sediments by transporting electrons over centimetre-scale distances to reduce oxygen. Here, we show that the freshwater cable bacterium *Electronema aureum* GS can respire insoluble electron acceptors under anoxic conditions via a versatile extracellular electron transfer (EET) system, supporting growth rates comparable to those under aerobic conditions. Using electrochemical and molecular biology analyses, we demonstrate that *E. aureum* GS engages in both direct and mediated electron transfer to electrodes, including at +600 mV vs. Ag/AgCl—an unusually high redox potential typically not accessed by electroactive bacteria. Two distinct cell-surface redox components were identified, which are metal-dependent, pH-sensitive, and heat-labile, consistent with outer-membrane-localised cytochromes. Moreover, the redox shuttle riboflavin accumulated extracellularly and enhanced current production in bioelectrochemical systems, indicating a role for soluble mediators in cable bacteria. Together, these findings reveal a previously unrecognised respiratory flexibility in *E. aureum* GS and highlight EET as a key alternative strategy for energy conservation in fluctuating redox environments.

## Introduction

Microbial life in sediment environments often faces the challenge of spatially separated electron donors and acceptors, as their availability is governed by dynamic chemical gradients^1^. For instance, although oxygen is the most thermodynamically favourable oxidant for hydrogen sulphide, sulphide produced in anoxic sediment layers can be effectively oxidised by biotic or abiotic processes that reduce nitrate or mineral oxides before diffusing into the oxic zone near the sediment surface ^2,3^. To overcome such limitations, cable bacteria have evolved specialized mechanisms for long-distance electron transfer, allowing them to sustain metabolic activity across steep redox gradients ^4,5^. Cable bacteria are filamentous members of the genera *Electronema* and *Electrothrix* that couple sulphide oxidation in deeper anoxic sediments to oxygen reduction at the surface by transporting electrons over centimetre distances^4,6^.

This unusual metabolism in cable bacteria electrically bridging redox zones has redefined our understanding of microbial respiration and sediment biogeochemistry. Cable bacteria function as living biological wires, forming electrical connections that shape redox zonation^7^, nutrient cycling^8^, pH profiles in sediments, and microbial interactions^9^. Beyond their role in sediment biogeochemistry, cable bacteria have been implicated in several ecological roles that have applied significance, including mitigation of methane emissions from rice-vegetated soils ^10^, degradation of organic pollutants in wastewater systems^11^, and protection of aquatic plants from sulphide toxicity in eutrophic environments^8^.

Cable bacteria exhibit high oxygen consumption rates (up to 2200 nmol O_2_ per mg protein per minute), suggesting oxygen as a strongly favourable electron acceptor^12^. This form of aerobic metabolism poses a fundamental challenge to cable bacteria, which live in environments where oxygen is temporally or spatially unavailable. Redox gradients in the sediment fluctuate due to seasonal shifts or bioturbation, often leading to conditions where cable bacteria filaments may encounter anoxia for prolonged periods of time^13,14,15^. In such conditions, alternative strategies for electron disposal are vital for survival. Some species of cable bacteria were found to reduce nitrate as an alternative electron acceptor^16^. However, growth rates under nitrate-reducing conditions have not been directly quantified.

Manganese and/or Fe-(oxyhydr)oxides are typically abundant in oxidized sediment layers and depending on their chemical speciation and crystalline structure, may persist in anoxic sediment^17^. These oxides could serve as alternative electron acceptors^18^ for cable bacteria during periods of anoxia, assuming that they can engage in extracellular electron transfer (EET) to solid-phase mineral oxides. However, cable bacteria remain uncultivated in pure culture, and their exclusive growth in sediments complicates efforts to isolate and characterize mineral reduction under controlled conditions. As a result, direct evidence for mineral-based EET in cable bacteria has remained elusive.

To circumvent this limitation, electrodes offer a stable, controllable, and quantifiable proxy for studying EET. By mimicking mineral electron sinks, electrodes enable the investigation of microbial redox activity in complex consortia ^19^. Critically, however, it has not been established whether cable bacteria can actively conserve energy and grow by respiring extracellular insoluble electron acceptors under anoxic conditions — a defining trait of classical electroactive bacteria such as *Geobacter* and *Shewanella* ^*20,21*^. While recent studies have reported current generation in bioelectrochemical systems (BESs) inoculated with cable bacteria-enriched sediments^22,23^, these studies did not conclusively demonstrate exclusive EET behaviour attributable to cable bacteria. Other studies involving direct electrochemical characterization using voltammetry revealed redox signals associated with cable bacteria filaments, hinting at the presence of electroactive components^24–26^. However, it remains unclear whether these signals arise solely from direct electron exchange at the cell surface or also involve internal conductive structures within the filaments. Thus, despite growing evidence that cable bacteria influence electrochemical activity in sediments, it has not been established whether cable bacteria themselves can perform EET to insoluble electron acceptors for energy conservation. In this study, we specifically aimed to determine whether cable bacteria can perform EET to insoluble electron acceptors under anoxic conditions for energy conservation and growth.

To investigate the mechanisms and physiological relevance of EET in cable bacteria, we studied the freshwater species *Electronema aureum* GS using a combination of electrochemical and molecular biology analyses. We found that *E. aureum* GS is capable of both direct and mediated EET to polarized electrodes, including at +600 mV vs. Ag/AgCl—an unusually high redox potential not typically accessed by conventional electroactive bacteria. Electrochemical profiling revealed two distinct redox peaks that are metal-dependent, and pH- and heat-sensitive, consistent with outer-membrane cytochromes. In addition, riboflavin enhanced current production, suggesting a role for soluble redox mediators in electron transfer. These findings demonstrate that *E. aureum* GS possesses a versatile EET system that supports respiration to solid electron acceptors under anoxic conditions and provide a basis for understanding how cable bacteria may conserve energy in the absence of soluble electron acceptors like oxygen and nitrate. Together, these findings expand the metabolic capabilities of cable bacteria, placing them among a growing class of microorganisms like *Escherichia coli*^*27*^, *Aeromonas hydrophila*^*28*^ and *Vibrio natriegens*^*29,30*^ that possess EET machinery despite well-known aerobic/fermentative metabolisms. This previously unrecognized EET capacity highlights alternative mechanisms for energy generation beyond canonical oxygen and nitrate reduction as the most favourable form of respiration and has implications for modelling of nutrient cycling mediated by cable bacteria.

## Results

### Redox-active metalloprotein centres on the cell surface of Electronema aureum GS

To investigate the potential for EET, differential pulse voltammetry (DPV) was applied to a sample of collected cable bacteria that migrated out of the sediment. For this aim, trench slides were used – a custom glass slide with a central cavity filled with cable bacteria-enriched sediment that facilitates the migration of intact filaments from the trench^6^. After incubating the trench slide for 48 hours, intact *E. aureum* GS filaments were isolated from the trench slide and deposited onto a carbon felt working electrode in a three-electrode system. DPV revealed two well-defined redox peaks around +250 mV and +600 mV versus Ag/AgCl (Fig. 1a), indicating the presence of active redox centres at these potentials. In contrast, the control, which consisted of the microbial community picked around *E. aureum* GS (while avoiding the filaments) – showed no such redox activity. This indicates that the observed electrochemical signals were intrinsic to *E. aureum* GS and not due to other bacteria in the enrichment.

**Figure 1.**
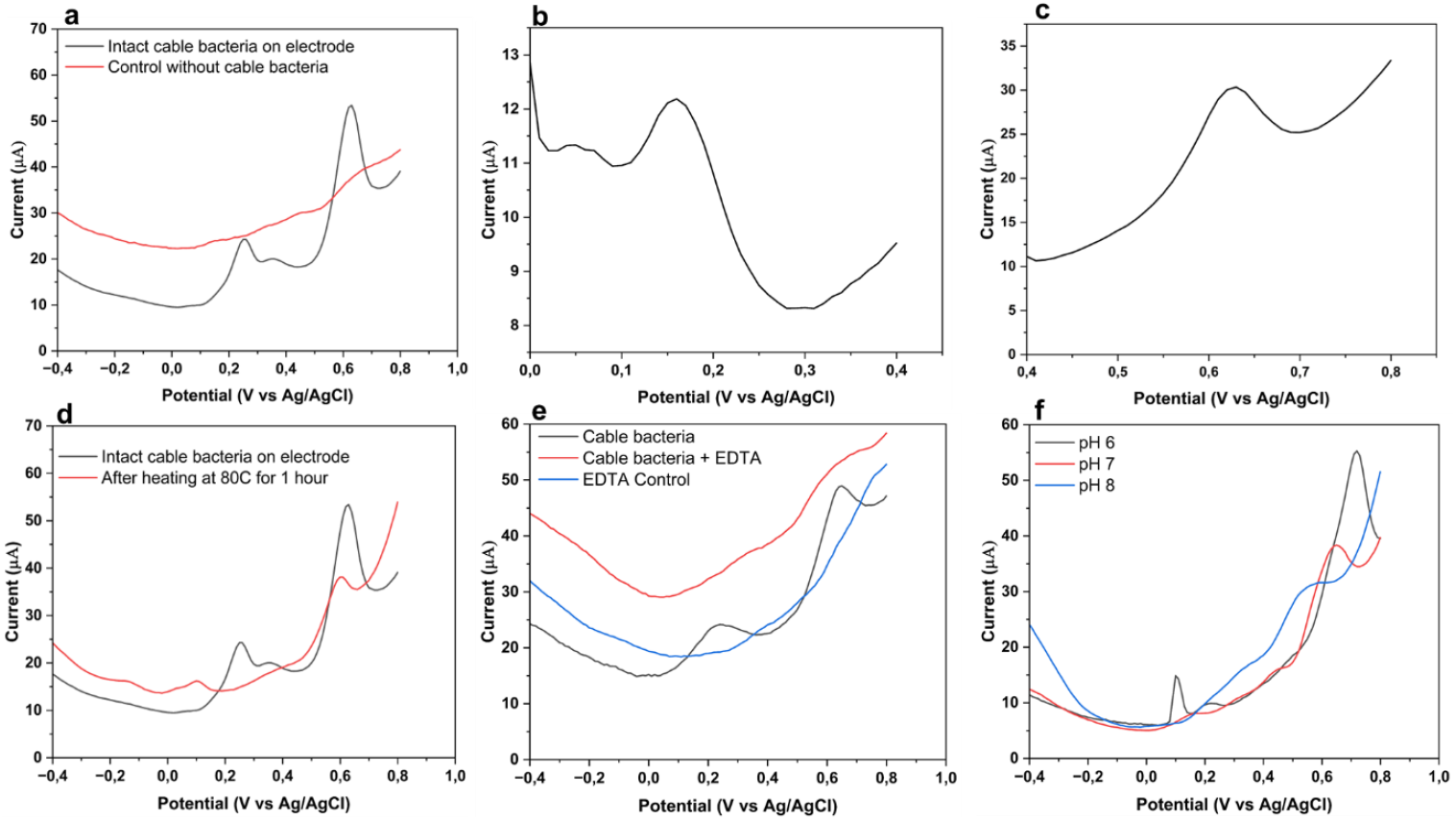
Electrochemical nature of cable bacteria’s cell-surface redox centres. a. Differential pulse voltammetry (DPV) of intact cable bacteria filaments deposited on carbon felt electrodes shows two distinct redox peaks at +250 mV and +600 mV vs. Ag/AgCl. No redox peaks are observed in controls containing other bacteria but lacking cable bacteria, indicating the redox features are unique to cable bacteria. b, c. DPV focused on narrow potential windows confirms the +250 mV and +600 mV vs. Ag/AgCl redox peaks are independently resolved. Together, these data suggest the presence of at least two distinct cell-surface redox centres in cable bacteria. d. DPV of cable bacteria before and after heating at 80°C for 1 hour. Heat treatment distorts both redox peaks, indicating thermal lability. e. Treatment of cable bacteria on electrodes with 50 mM EDTA destroyed both redox peaks, indicating that metal cofactors are required for redox activity. f. DPV of cable bacteria on electrodes at pH 6, 7, and 8. The redox peaks shift with increasing pH, consistent with activity exhibited by redox-active outer-membrane proteins.

*E. aureum* GS filaments did not undergo any treatment before deposition on electrodes and remained hydrated, implying that the redox-active components must reside on cell surface of intact filaments. Sequential DPV scans across narrower potential windows independently resolved both redox peaks, indicating that the two signals arise from distinct and electrochemically independent moieties (Fig. 1b, 1c). Supporting this, DPV scans within the same potential ranges showed bidirectional redox behaviour for both peaks (Fig. S1), consistent with electron transfer mediated by redox-active proteins. This suggests that *E. aureum* GS contains at least two outer-membrane redox centres that could facilitate electron exchange with extracellular electron acceptors.

To examine the biochemical nature of the cell-surface redox-active components, we subjected cable-bacteria-coated electrodes to heat treatment and chelation. Heating the samples to 80°C for one hour substantially distorted both redox peaks (Fig. 1d), indicating thermal denaturation of the redox agents, consistent with the redox agents being protein-based. Similarly, chelation with 50 mM ethylenediaminetetraacetic acid (EDTA) destroyed both redox peaks (Fig. 1e), implicating the presence of essential metal cofactors, consistent with redox-active outer-membrane metalloproteins. We also tested the effect of sodium azide, a known inhibitor of heme-containing redox enzymes. Upon addition of 10 mM sodium azide, both the +250 mV and +600 mV peaks were abolished (Fig. S2), consistent with the inhibition of outer-membrane metalloproteins such as cytochromes or terminal oxidases^31–33^. Control experiments with azide alone showed no intrinsic redox peaks, confirming that the loss of signal was specific to inhibition of cable bacteria redox activity.

The redox potentials exhibited consistent negative shifts with increasing pH across the range 6-8 (Fig. 1f), consistent with proton-coupled electron transfer mechanisms typical of outer-membrane cytochromes^34,35^. Together, these data strongly support the presence of two outer-membrane, proteinaceous, metal-cofactor-containing redox centres in *E. aureum* GS, capable of direct electron exchange with extracellular electron acceptors.

### Electronema aureum GS performs EET under anoxic conditions

To test whether these two redox centres could be implicated in EET under physiological conditions and have a metabolic role, we inoculated sediment enrichments of *E. aureum* GS^36^ into three-electrode BESs with working electrodes poised at +250 mV and +600 mV vs. Ag/AgCl. In both cases, *E. aureum* GS-inoculated sediments produced substantial anodic currents compared to control sediments lacking *E. aureum* GS (Fig. 2a, 2b). At +250 mV, the maximum current was higher than at +600 mV, suggesting a more energetically favourable EET pathway at this potential. Interestingly, *E. aureum* GS-containing BESs poised at +600 mV (vs Ag/AgCl) exhibited a steady current over time. Control BESs at +600 mV, which contained only the associated microbial community but not *E. aureum* GS filaments, produced negligible current. We also tested a pure culture of *Shewanella oneidensis* for its EET capability at +600mV (Fig. S3). *S. oneidensis* failed to generate appreciable current at +600 mV under identical conditions, further reinforcing the high-potential EET ability of cable bacteria. This distinct response underscores the unique capacity of *E. aureum* GS to utilize unusually high-potential outer-membrane proteins.

**Figure 2.**
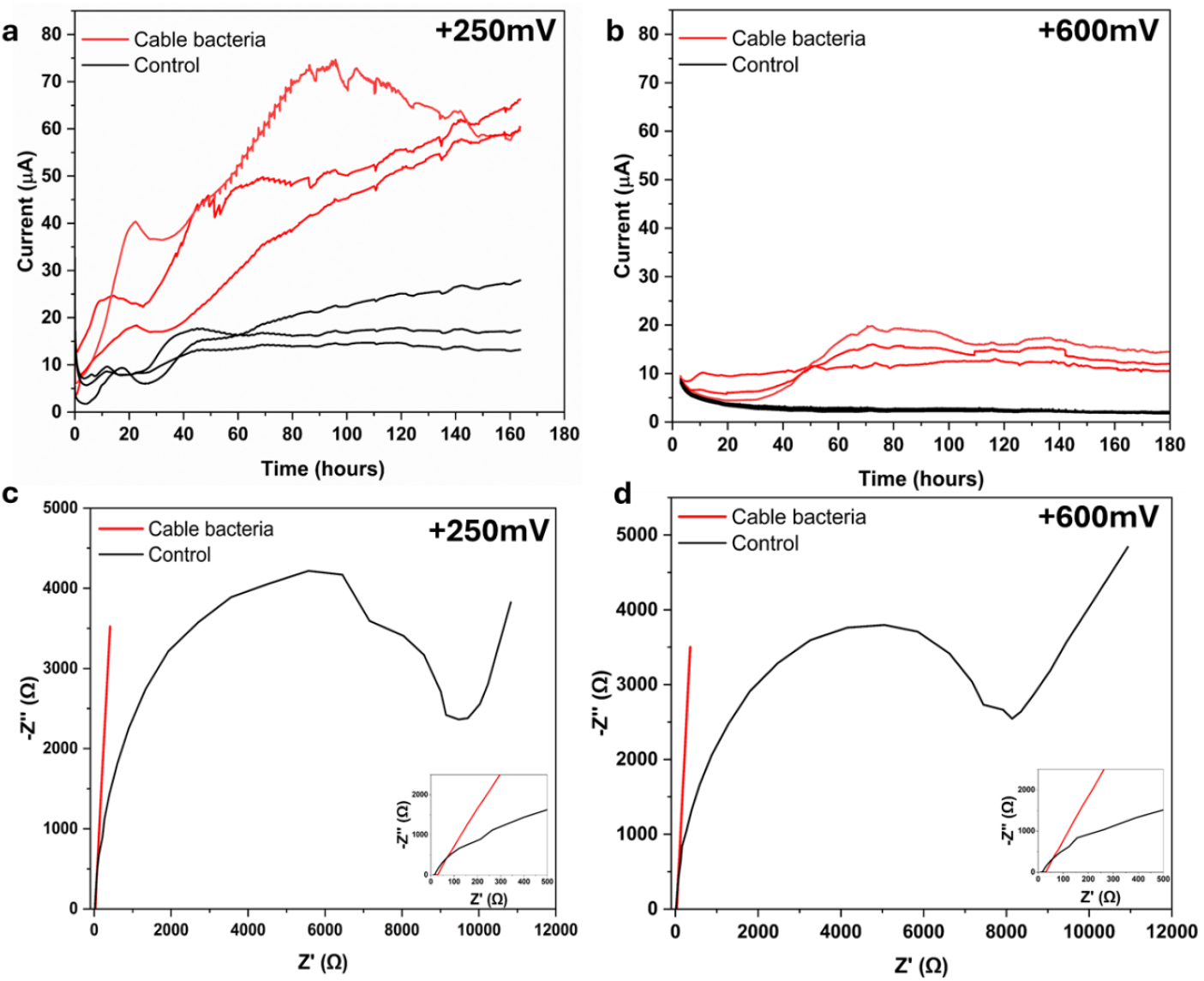
Electrode respiration by cable bacteria in sediments under anoxic conditions. a. Chronoamperometry of sediment BES poised at +250 mV vs. Ag/AgCl. Sediments containing cable bacteria generate significantly higher currents than cable bacteria-free controls. Triplicates are presented. b. Similar experiment at +600 mV shows cable bacteria-containing sediments still produce elevated current, though lower than at +250 mV. Triplicates are presented. These data demonstrate that cable bacteria can perform anoxic electrode respiration at two different redox potentials. c, d. Electrochemical impedance spectroscopy (EIS) of cable bacteria and control at +250 mV and +600 mV vs. Ag/AgCl. At both potentials, cable bacteria exhibited significantly lower charge transfer resistance. The controls display high impedance. Insets show magnified views of the low-impedance regions (Z’ < 500 Ω).

To confirm that the observed redox activity at +600 mV was not due to electrochemical oxygen production, we performed control experiments in BESs with phosphate buffer as the electrolyte. Oxygen generation was only detectable at +1.2 V vs. Ag/AgCl. Using the Nernst equation, we calculated the theoretical equilibrium oxygen concentration at +0.6 V to be approximately 5.27 × 10−^24^ µM—an exceedingly low and physiologically irrelevant value (Section 1, Supplementary Information). These findings demonstrate that oxygen evolution at +0.6 V is thermodynamically negligible, and the redox activity observed in *E. aureum* GS at this potential reflects biological electron transfer to the electrode, rather than oxygen-dependent respiration.

Microscopy of trench slides made from these BESs at the end of chronoamperometry experiments confirmed the continued presence of morphologically intact, motile, living cable bacteria filaments (Fig. S4). This indicates that cable bacteria can survive without oxygen, possibly substituting oxygen respiration for electrode respiration.

To further investigate the electron transfer capabilities of *E. aureum* GS at the electrode interface, we performed electrochemical impedance spectroscopy on carbon felt electrodes with intact *E. aureum* GS filaments. Control electrodes were prepared from the same sediment enrichment but excluded *E. aureum* GS, retaining only other bacteria. Nyquist plots revealed distinct impedance responses for *E. aureum* GS and the control (Fig. 2c, d). *E. aureum* GS displayed substantially lower impedance in the high-frequency region compared to controls at both potentials, indicating reduced charge transfer resistance and more efficient interfacial electron transfer.

The low impedance at +250 mV correlates well with the higher current generation and growth rates observed in chronoamperometry. Notably, reduced impedance was also observed for *E. aureum* GS at +600 mV, supporting the unique ability of *E. aureum* GS to respire to high-potential electron acceptors. These impedance results support the pH-dependent, EDTA and azide-sensitive redox peaks observed in DPV, which implicate outer-membrane cytochromes in electron transfer. While EIS does not resolve individual redox transitions, the consistently reduced charge transfer at both potentials indicates that these redox systems are electrochemically competent and contribute to EET. Together, the data support a model in which *E. aureum* GS employs multiple outer membrane redox components to support EET across a broad potential window.

### Cable bacteria engage in mediated EET via soluble redox shuttles

We then investigated whether *E. aureum* GS could engage in mediated electron transfer. We constructed trench slide BESs with a built-in carbon fibre working electrode and a chlorinated silver wire pseudoreference electrode. Electrodes were poised at either +250 mV or +600 mV vs. Ag/AgCl, with unpoised electrodes serving as controls. In trench slide BESs inoculated with *E. aureum* GS, we added water spiked with 5 mM riboflavin, a known redox mediator (Fig. 3a). Chronoamperometry at +250 mV showed that cable bacteria-containing inocula generated higher currents in the presence of riboflavin than without it (Fig. 3b). Cable bacteria in the absence of externally added riboflavin still produced appreciable current, except it was lower, while cable bacteria-free sediments produced minimal current under both conditions. This hierarchy supports the ability of cable bacteria to use both direct and mediator-based EET pathways.

**Figure 3.**
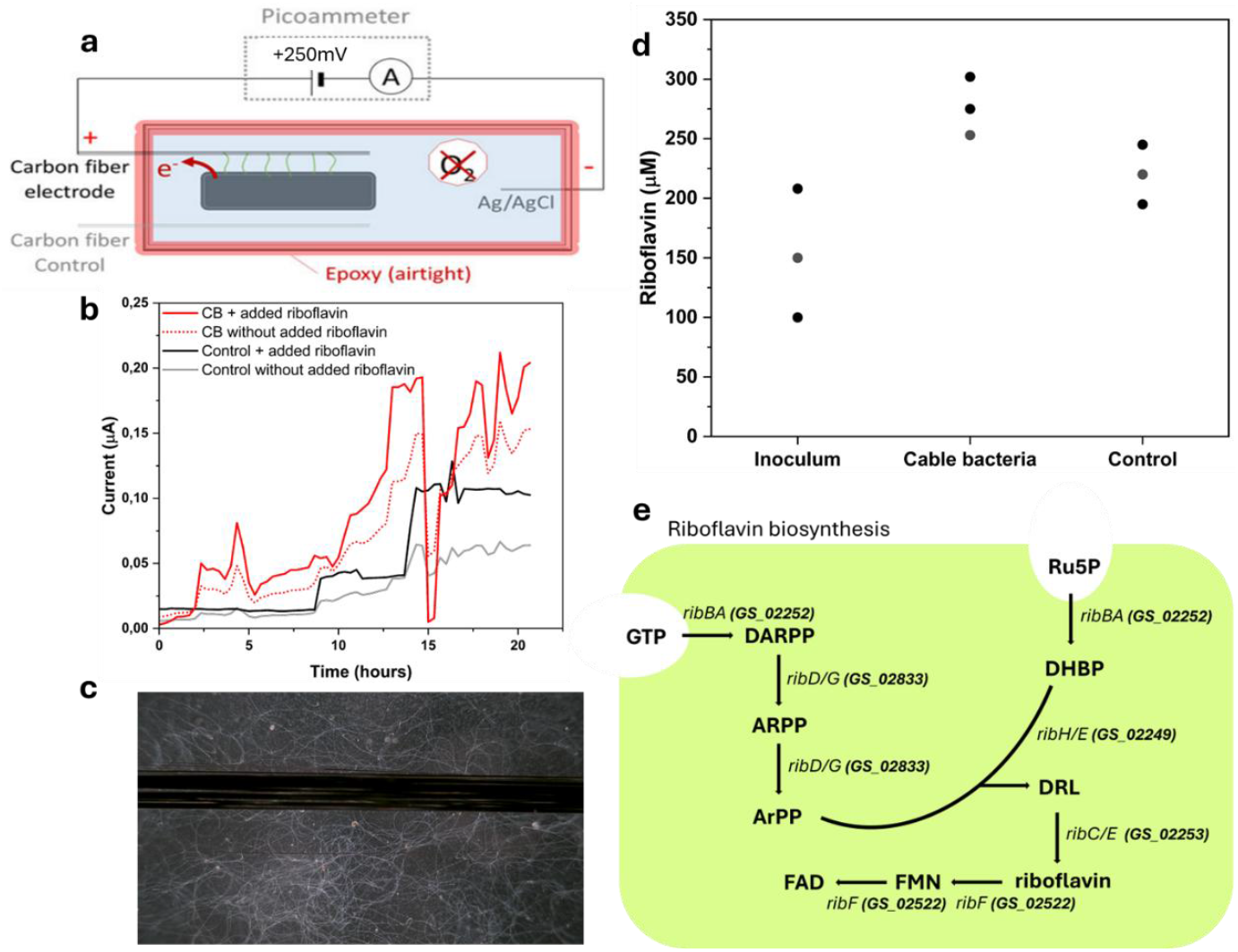
Evidence for mediated electron transfer in cable bacteria. a. Schematic of trench-slide BES incorporating a carbon fibre working electrode and chlorinated silver pseudoreference electrode. b. Chronoamperometry in this setup with the electrode poised at +250 mV shows that cable bacteria-containing inocula produce higher currents in the presence of externally spiked riboflavin (mean data shown). c. Light microscopy images of cable bacteria filaments clustering at the poised electrode surface. d. Plot showing riboflavin concentrations (µM) measured in three different sample types: inoculum (native sediment), BESs containing cable bacteria, and control. Cable bacteria enrichments exhibited higher riboflavin levels compared to controls and inoculum, consistent with flavin-mediated extracellular electron transfer. e. Schematic overview of the main genes that play a role in the riboflavin biosynthesis in *Electronema aureum* GS. **Genes:** ribF, bifunctional flavokinase/FAD synthetase; ribC/E, riboflavin synthase; ribBA, bifunctional enzyme GTP cyclohydrolase II/3,4− DHBP synthase; ribD/G, bifunctional deaminase/reductase; ribH/E, lumazine synthase. **Metabolites:** FAD, flavin adenine dinucleotide; FMN, flavin mononucleotide; DRL, 6,7− dimethyl−8−ribityllumazine; DARPP, 2,5−diamino−6−ribosyl−amino−4(3H)pyrimidinedione 5′− phosphate; ARPP, 5−amino−6−ribosyl−amino−2,4(1H,3H)pyrimidinedione 5′−phosphate; ArPP, 5−amino−6−ribityl−amino−2,4(1H,3H)pyrimidinedione 5′−phosphate; ArP, 5−amino−6−ribityl− amino−2,4(1H,3H)pyrimidinedione; DHBP, 3,4−dihydroxy−2−butanone−4−phosphate; GTP, guanosine 5′−triphosphate; Ru5P, ribulose 5−phosphate. **Uniprot accession numbers**: A0A521G3Y1 (*GS_02249*); A0A521G3Y6 (*GS_02252*); A0A521G3X3 (*GS_02253*); A0A521FYE6 (*GS_02522*); A0A521G0P9 (*GS_02833*)

Live microscopy of the trench slide BES revealed cable bacteria filaments actively aggregating near the electrode surface (Fig. 3c), suggestive of electrotaxis, a phenomenon previously observed in cable bacteria^23^ and in other electroactive microbes^37^. HPLC-based quantification of riboflavin from regular BESs indicated that cable bacteria treatments accumulated extracellular riboflavin at higher concentrations compared to controls (Fig. 3d), suggesting that cable bacteria may either produce or accumulate redox mediators.

In support of the electrochemical and biochemical evidence for flavin-mediated electron transfer, we further investigated the genomic potential of *E. aureum* GS for flavin biosynthesis and export. Bioinformatic analysis confirmed the presence of key genes involved in the riboflavin biosynthesis pathway, including *ribBA, ribC/E, ribD/G, ribH/E*, and *ribF* (Fig. 3e), indicating that *E. aureum* GS possesses the complete enzymatic machinery to endogenously produce flavin derivatives such as FMN and FAD. In addition, structural homology search using FoldSeek for *E*.*aureum* GS AlphaFold models identified a membrane protein (GS_00133) to have the closest structural similarity to both known flavin exporters Bfe and YeeO from *Shewanella oneidensis* and *Escherichia coli*, respectively (Fig. S5). The predicted membrane localization of GS_00133 is based on the presence of 12 transmembrane helices determined by DeepTMHMM. However, the closest structural homolog of GS_00133 in Alphafold protein structure database is Lipid III flipase, which is inner membrane transporter involved in cell surface biosynthesis. These findings suggest that *E. aureum* GS should be able to to synthesize flavins *de novo*.

### Electronema aureum GS uses EET to grow under anoxia

To determine whether the observed EET supports energy conservation and growth of *E. aureum* GS, we performed qPCR targeting its 16S rRNA gene on sediment samples recovered from trench slide BESs. Across all replicates, sediments from +250 mV poised electrodes showed a marked increase—up to two orders of magnitude—in *E. aureum* GS 16S rRNA gene copy numbers compared to unpoised controls and initial inocula (Fig. 4). Although the numbers at +600 mV were comparatively lower, they were higher relative to controls, indicating survival of *E. aureum* GS even at higher redox potentials. The qPCR data align closely with the observed chronoamperometric current profiles, indicating that *E. aureum* GS is selectively enriched and can grow via EET towards poised electrodes.

**Figure 4.**
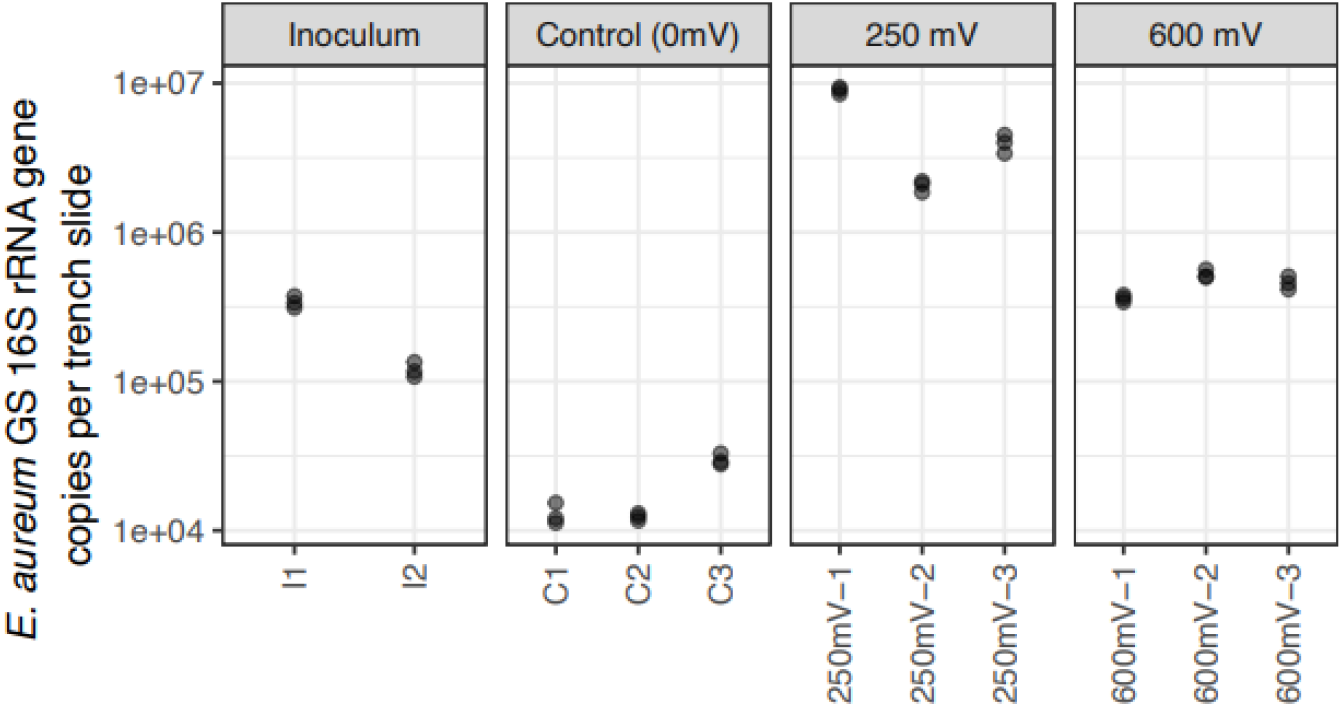
Quantification of *E. aureum* GS abundance on poised electrodes via qPCR. Plot showing 16S rRNA gene copy number per gram of sediment for inoculum (I1, I2) unpoised controls (C1–C3) and replicate BESs poised at +250 mV and +600 mV vs. Ag/AgCl. Triplicates are presented. Enrichment of *E. aureum* GS is especially pronounced at + 250 mV.

Overall, these findings demonstrate that electrodes poised at +250 mV support active extracellular electron transfer by *E. aureum* GS under anoxic conditions, enabling energy conservation and growth comparable to aerobic conditions.

We estimated doubling times for *E. aureum* GS based on 16S rRNA gene abundance measured by qPCR. Sediment samples of equal mass (0.3 g) were weighed for DNA extraction across all conditions. Assuming exponential growth over the 3.5-day incubation period (84 hours), we calculated the number of generations (n) as:

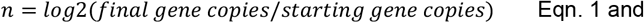

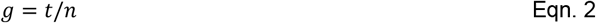

Where g is the doubling time and t = 84 hours. For electrodes poised at +250 mV, the calculated doubling times ranged from 13.5 to 36.8 hours, depending on the minimum and maximum gene copy numbers used (section 2, supplementary information provides calculations). Using mean values across replicates, the average doubling time was approximately 20 hours. These estimates align closely with doubling times reported for *Electrothrix* species under oxic conditions (e.g., <20 h) ^38,39^, indicating that poised electrodes at +250 mV can support cable bacteria growth rates comparable to oxygen respiration.

In contrast, for BESs poised at +600 mV, estimated doubling times were substantially longer. Even in the best-case scenario, the doubling time was at least 39.4 hours, and in some comparisons, gene copy numbers decreased or barely changed relative to the inoculum, suggesting negligible or no net growth. Mean-based estimates yielded a doubling time of approximately 115.5 hours at +600 mV. These findings indicate that while cable bacteria can sustain EET and remain metabolically active at high redox potentials, growth is considerably slower than at +250 mV.

To examine the microbial community in sediment BESs enriched during chronoamperometry, we performed 16S rRNA gene amplicon sequencing of the microbial communities in BESs poised at +250 mV and +600 mV. In BES poised at +250 mV, *E. aureum* GS was highly enriched compared to control inocula (Fig. S6a). BES poised at +600 mV also showed *E. aureum* GS enrichment (Fig. S6b), though to a lesser degree, suggesting a more selective or energetically costly EET pathway at this higher potential. These results align well with chronoamperometry and qPCR data, suggesting the involvement of *E. aureum* GS in EET.

## Discussion

Previous electrochemical studies hinted at redox-active components in cable bacteria, but their physiological roles remained unresolved. Meysman et al^26^ previously hypothesised the presence of cytochromes involved in EET based on voltammetry experiments. Other studies reported attraction of cable bacteria to electrodes poised at oxidising potentials (200-300mV vs Ag/AgCl) suggesting EET activity^22,23,40^. However, these studies did not conclusively demonstrate the EET activity attributable to cable bacteria and did not provide mechanistic details.

In contrast, our study provides direct physiological evidence that *E. aureum* GS performs EET through both direct and mediated pathways, and that this process supports growth under anoxic conditions. Despite the absence of pure culture isolates, our sediment-based BESs provided a clear and direct attribution of EET activity to cable bacteria. This extends the current understanding of cable bacteria metabolism, which has primarily centred on long-distance electron transport enabling sulphide oxidation coupled to oxygen or nitrate reduction. Here, we show that cable bacteria can also reduce insoluble extracellular electron acceptors such as electrodes—broadening their metabolic repertoire. Differential pulse voltammetry identified two distinct redox centres (∼+250 mV and ∼+600 mV vs. Ag/AgCl), consistent with outer-membrane cytochromes based on their pH sensitivity and inactivation by heat, azide and chelators. These surface-localised redox components reflect EET mechanisms seen in model electroactive bacteria ^34,35^ and support emerging evidence that such systems are widespread among sediment microbes capable of extracellular respiration ^41,42^.

Recent comparative genomic and structural analysis of different cable bacteria^5^ provides additional context to the outer-membrane redox centres identified here. Several extracytoplasmic cytochromes conserved across available cable bacteria genomes display homology to known EET proteins in model bacteria, including multiheme cytochromes from *Shewanella* and *Geobacter* ^5,43^. These homologs retain key features such as β-barrel domains and multiple heme-binding CXXCH motifs typical of electron-transferring cytochromes^44^.

A particularly notable feature of cable bacteria EET metabolism is their ability to transfer electrons to electrodes poised at +600 mV vs. Ag/AgCl—an unusually high potential beyond the range of model electroactive bacteria such as *S. oneidensis* and *Geobacter sulfurreducens*. Sustained current generation at this high potential suggests that *E. aureum* GS is capable of respiration to high-potential electron acceptors. While the exact physiological role of the redox centre at +600 mV remains to be fully elucidated, it is plausible that it is associated with oxygen reduction. This is particularly relevant given that most cable bacteria, including *E. aureum* GS, lack genes encoding canonical cytochrome c oxidases^43^, implying the presence of an alternative, yet unidentified, high-potential terminal oxidase system. The redox activity observed at +600 mV may therefore represent a component of this non-canonical oxygen reduction pathway.

In addition to direct EET, our data reveal a role for MET via soluble redox shuttles. Riboflavin supplementation significantly enhanced current production, and riboflavin was found to accumulate in cable bacteria-containing systems. These findings suggest that cable bacteria not only respond to exogenous mediators but may also produce or accumulate riboflavin to support MET with other community members. Such dual-mode electron transfer – direct via redox-active proteins and mediated via soluble shuttles – has been observed in *S. oneidensis*^*21*^ and *Aeromonas hydrophila*^*28,45*^ among other electroactive bacteria but rarely demonstrated in uncultivated organisms. Our results provide one of the few examples, alongside recent studies such in *E. coli* ^27^, where EET strategies are deployed in organisms despite fermentative and aerobic respiratory capabilities. While bioinformatics analyses detected protein homologues of Bfe and YeeO, further experimental evidence is needed to confirm the activity of these exporters.

In addition, we suggest that the same EET mechanisms may be involved in syntrophic interactions with flocking microorganisms^9,42,46^. This is analogous to mechanisms observed in anaerobic methane-oxidizing (ANME) consortia, where archaea donate electrons directly to partnering bacteria that use different terminal electron acceptors^47,48^. In the case of cable bacteria, our findings raise the possibility that flocking microbes may provide electrons via direct interspecies electron transfer, allowing cable bacteria to access reduced substrates such as sulphide or organics in the absence of regular electron donors. Such interactions could expand the ecological niche of cable bacteria and support their persistence in otherwise electron-donor-limited environments. The bidirectionality of the redox peaks raises the intriguing possibility that cable bacteria are capable of both electron donation and uptake, opening up the exploration for electrotrophic growth in future studies. These findings also have implications for nutrient cycling in both natural biogeochemical processes and engineered systems, where cable bacteria may engage in interactions with minerals and participate in metal mobilisation.

## Methods

### Trench slide preparation

Trench slides were prepared to obtain intact filaments of *Electronema aureum* GS for electrochemical analysis. The trench slide consisted of a standard glass microscope slide into which a central rectangular cavity (∼1 mm deep) was etched, as previously described ^6^. This cavity served as an environment for the inoculation of cable bacteria-containing sediment.

The trench was filled with sediment enriched with a single strain of *E. aureum* GS ^36^. Sterile water was gently added over the sediment, and a thin glass coverslip was placed carefully on top to maintain humidity and prevent desiccation. The slide was incubated horizontally for 48 hours in a sealed humid chamber at room temperature to allow cable bacteria filaments to migrate from the trench towards the edges of the trench.

Following incubation, the coverslip was removed by carefully adding water along one edge of the coverslip and gently sliding the coverslip aside without disturbing the underlying cable bacteria filaments. Using a stereomicroscope, intact cable bacteria filaments extending from the trench were visually located and selectively pipetted (Check https://doi.org/10.6084/m9.figshare.30093571.v1 for videos demonstrating this). For the control, material was pipetted from a region of the same trench slide that was free of visible cable bacteria but contained other microbial community members.

### Differential pulse voltammetry (DPV) and electrochemical impedance spectroscopy (EIS)

The isolated cable bacteria filaments from trench slides were deposited directly onto carbon felt electrodes serving as the working electrodes in three-electrode bioelectrochemical cells (30 ml working volume). An Ag/AgCl reference electrode and a platinum wire counter electrode completed the configuration. Phosphate-buffered saline (PBS) (pH 7.4) was used as the electrolyte. DPV was conducted in a scan range of –800 mV to +800 mV vs. Ag/AgCl, using a pulse amplitude of 50 mV, a step size of 1 mV, a pulse duration of 0.3 s, and a pulse interval of 1 s. EIS was done in potentiostatic mode based on the peaks obtained in DPV (+250mV and +600mV). The potential amplitude was 30 mV. All electrochemical measurements were performed using a MultiPalmSens 10-channel potentiostat (PalmSens BV, Netherlands) operated with MultiSens software.

### Heating, chelation, pH and azide experiments on cable bacteria redox activity

To assess the biochemical nature of the redox-active components in *E. aureum* GS, intact filaments were collected from trench slides as described above and deposited onto carbon felt working electrodes within three-electrode bioelectrochemical systems (BES). Initial DPV was performed to establish baseline electrochemical profiles.

For the thermal denaturation assay, the entire BES, containing cable bacteria-deposited electrodes, was placed in a water bath at 80 °C for 1 hour. After cooling to room temperature, the same DPV protocol was repeated to assess changes in redox activity post-heating.

In a separate set of experiments, chelation was used to evaluate the involvement of metal cofactors. Following initial DPV scans of cable bacteria-deposited electrodes, 50 mM EDTA was added directly to the electrolyte, and the system was incubated for 15 minutes at room temperature. Subsequent DPV measurements were conducted without removing the sample to determine the effect of metal chelation on redox peak behaviour.

To assess the pH sensitivity of the observed redox features, additional DPV measurements were performed in buffered electrolytes adjusted to pH 6.0, 7.0, and 8.0 using PBS titrated with NaOH or HCl as needed. For each pH condition, fresh cable bacteria filaments were collected and deposited onto new carbon felt electrodes to avoid confounding effects from prior treatments. All electrochemical parameters were consistent with the protocol described previously. To test the susceptibility of the redox components to azide inhibition, 10 mM sodium azide was added to the BESs after initial DPV measurements. The system was incubated for 15 minutes at room temperature before repeating DPV scans under identical conditions.

### Chronoamperometry in sediment bioelectrochemical systems containing cable bacteria

To evaluate the capacity of *E. aureum* GS to perform extracellular electron transfer under anoxic conditions, sediment-based BESs were constructed for chronoamperometry. Single-strain enriched sediment cultures of *E. aureum* GS were used as the inoculum. For each BES, approximately 10 mL of enriched sediment was transferred into sterile 30 mL anaerobic serum bottles, which served as the electrochemical reactors.

Carbon felt electrodes were used as the working electrodes and were embedded within the sediment matrix to facilitate contact with active cable bacteria. An Ag/AgCl reference electrode and a platinum wire counter electrode were positioned in the overlying anoxic water layer. The headspace was flushed with N_2_ for 20 minutes to ensure anoxic conditions.

Chronoamperometry was performed using a MultiPalmSens 10-channel potentiostat (PalmSens BV) by poising the working electrode at either +250 mV or +600 mV vs. Ag/AgCl. Current was recorded continuously over time to monitor the development of bioelectrochemical activity. Control BESs were identically assembled using sediment containing associated microbial communities but not cable bacteria filaments. All experiments were conducted in biological triplicates, and trench slides were done after the chronoamperometry experiments to confirm the presence or absence of cable bacteria filaments.

### 16S rRNA gene sequencing

16S rRNA gene sequencing was performed after chronoamperometry experiments to determine the composition of the microbial community from the three-electrode cells. DNA was extracted from each bioelectrochemical system, and the V3–V4 hypervariable region of the 16S rRNA gene was amplified using primers Bac341F (5′-GGGCATCATGATGCGCCTG-3′) and Bac805R (5′-TCRACCTTCTCGCACTTCCA-3′)^49^ at a final concentration of 0.2 pmol/μL. PCR amplification was performed using 2× KAPA HiFi HotStart ReadyMix (Roche, Switzerland) in 25 μL reactions containing 2.5 μL template DNA. The thermal profile included an initial denaturation at 95 °C for 15 min, followed by 30 cycles of 95 °C for 30 s, 54 °C for 30 s, and 72 °C for 20 s, with a final extension at 72 °C for 5 min.

A second PCR using primers containing Illumina adapter overhangs (Bac341F-adapt: 5′-TCGTCGGCAGCGTCAGATGTGTATAAGAGACAGCCTACGGGNGGCWGCAG; Bac805R-adapt: 5′-GTCTCGTGGGCTCGGAGATGTGTATAAGAGACAGGACTACHVGGGTATCTAATCC) was performed for 15 cycles under the same conditions. Indexing PCR was conducted using Nextera XT Index Kit primers (Illumina) with 15 cycles and an annealing temperature of 55 °C.

Sequencing libraries were purified and pooled for paired-end sequencing on an Illumina MiSeq platform using the V3 chemistry. Raw reads were processed using *cutadapt* v3.5 to remove primer sequences^50^. Amplicon sequence variants (ASVs) were inferred using *DADA2* v1.28.0^51^ in R v4.5.1. Forward and reverse reads were trimmed to 250 and 200 bp, respectively. Taxonomic classification was performed against the SILVA 138.1 SSU reference database ^52,53^.

#### Trench slide BES containing cable bacteria

A customized bioelectrochemical system was designed using a trench slide platform (Fig. S5). The trench slide was inoculated with sediment enriched with *E. aureum* GS. Control trench slides were inoculated with sediment without cable bacteria. Two parallel bundles of carbon fibre wires (10 µm diameter; T300SC, Toray, Japan) were affixed on either side of the trench at a 5 mm spacing. One bundle functioned as the working electrode, while the opposing bundle remained unpoised and served as a negative control. At a distance of 20 mm from the trench, a chlorinated silver wire was positioned to act as both the pseudo-reference and counter electrode. Following inoculation, the trench was overlaid with nitrogen-sparged distilled water with or without 5 mM riboflavin, and a coverslip was placed over the setup. The slide was incubated at 25 °C for 4 h to allow cable bacteria filaments to emerge from the sediment into the overlying water. After this period, the coverslip was sealed with epoxy resin (Loctite Instant Mix, Henkel, USA) to minimize oxygen intrusion. A potential of +200 mV was applied between the working electrode and the Ag wire using a picoammeter (Unisense, Denmark), and current was continuously monitored with SensorTrace software (Unisense).

#### qPCR

Quantification of *Electronema aureum* GS from trench slide BESs was performed using strain-specific primers targeting the 16S rRNA gene: GS68F (5′-GGTAGTTTCCTTCGGGGGAC-3′) and GS178R (5′-CTTCGGCAATGCGGCGTATC-3′).^13^

DNA was extracted from 0.5 g of sediment or electrode material using the DNeasy PowerLyzer PowerSoil Kit (Qiagen), following the manufacturer’s protocol. Each 20 μL qPCR reaction contained 2 μL of DNA extract, RealQ Plus 2× Master Mix Green (Ampliqon, Odense, Denmark), 0.1% BSA, and 0.5 pmol/μL of each primer. A synthetic DNA fragment (GenScript, Rijswijk, Netherlands) corresponding to positions 1–200 of the GS 16S rRNA gene was used to generate a standard curve.

Reactions were run on a Stratagene Mx3005P qPCR system (Agilent Technologies, Santa Clara, CA, USA) using the following thermal profile: 95 °C for 15 min, followed by 45 cycles of 95 °C for 30 s, 60 °C for 30 s, 72 °C for 20 s, and 95 °C for 15 s (data acquisition step). Melting curve analysis from 60 °C to 95 °C confirmed the specificity of amplification. Cell abundance was normalized per gram of sediment.

#### Bioinformatic methods for identification of riboflavin biosynthesis and export homologs

The proteome of *E. aureum* GS was searched for the presence of sequence homologs to riboflavin biosynthesis enzymes using Standart Protein BLAST for protein-protein searches ^54,55^. The AlphaFold database structure predictions AF-Q8EIX5-F1 (Bfe; *S. oneidensis* strain MR-1) and AF-A0A0H3EZQ8-F1 (YeeO; *E. coli* strain K12) were found through UniProt accession (The Uniprot Consortium, 2023) and run in FoldSeek^56^ against the AFDB50 database of *E. aureum* GS. Cellular localization of the potential flavine exporter GS_00133 was confirmed using TMHMM^57^.

## Acknowledgments

K.A. and M.L. acknowledge funding from the European Union’s Horizon Research and Innovation Program under the Marie Sklodowska-Curie grant agreement (project 101109777 – Cable Electricity O_2_ and project 101210107 — LiveWire respectively). L.D. acknowledges that a part of the work presented here was supported by the Carlsberg Foundation, grant CF24-0116. All authors acknowledge funding from the Danish National Research Foundation (Center for Electromicrobiology, DNRF136). The authors thank Dimitri Stamatis for his assistance with the HPLC, Lars Børregård Pedersen and Cecilie Sølund Bolø for help with trench slides and Lykke Bamdali for assistance with qPCR and sequencing.

## Notes

https://doi.org/10.6084/m9.figshare.30093571.v1

### Competing Interest Statement

The authors have declared no competing interest.

