## Supplementary Information for "Extracellular electron transfer in cable bacteria enables growth rates comparable to aerobic respiration"

### Section 1: Estimation of Oxygen Generation at +0.6 V vs. Ag/AgCl

The experimental setup consisted of a three-electrode cell (as described in the methods section) with phosphate buffer as the electrolyte. The electrode was poised at +0,6V vs Ag/AgCl.

To evaluate whether oxygen could be generated electrochemically at an applied potential of +0.6 V vs. Ag/AgCl, we used the Nernst equation for the half-reaction:

 O₂ + 4H⁺ + 4e⁻ → 2H₂O

The standard electrode potential for this reaction is:

 E°(O₂/H₂O) = +1.23 V vs. SHE

Given that the Ag/AgCl reference electrode (sat. KCl) is +0.197 V vs. SHE, the applied potential of +0.6 V vs. Ag/AgCl corresponds to:

 E_applied = 0.6 V + 0.197 V = 0.797 V vs. SHE

We apply the Nernst equation:

 E = E° - (RT/nF) * ln(1/[O₂])

Where:
 R = 8.314 J/mol·K
 T = 298.15 K
 F = 96485 C/mol
 n = 4

Substituting values:
 [O₂] = exp[-(4 × 96485) / (8.314 × 298.15) × (1.23 - 0.797)]
 = 5.27 × 10⁻^24^ µM

At an applied potential of +0.6 V vs. Ag/AgCl, the theoretical equilibrium concentration of molecular oxygen at the electrode surface is ~5.27 × 10⁻^24^ µM, an extremely low value that is many orders of magnitude below physiological or detectable levels. Oxygen production is thermodynamically negligible at +0.6 V, and cable bacteria activity under these conditions is unlikely to involve oxygen as an electron acceptor. This calculation confirms that the current observed at +0.6 V originates from alternative metabolic pathways such as extracellular electron transfer (EET), and not from oxygen-related processes.

#### Section 2: Doubling times of *E. aureum* GS under anoxic conditions

To estimate the doubling time of *Electronema aureum* GS in trench slide bioelectrochemical systems (BES), we used qPCR-derived 16S rRNA gene copy numbers measured in the inoculum and after 3.5 days (84 hours) of incubation on electrodes poised at +250 mV or +600 mV vs. Ag/AgCl. Equal sediment mass (~0.3 g) was used for DNA extraction across inoculum and endpoint samples. The following assumptions and formulae were used:

– Growth was exponential (log phase) over the 84-hour period.
– Gene copy number is assumed to be proportional to cell abundance.

– All reported values are total 16S rRNA gene copies per 0.3 g of wet sediment (i.e., the amount used for DNA extraction per sample).

Calculations

Number of generations:
 n = log2(N_final / N_start)
Doubling time:
 g = t / n, where t = 84 hours

##### +250 mV Condition

Start (inoculum) values:
 N_start, min = 119,600
 N_start, max = 421,333

Final values (after 3.5 days at +250 mV):
 N_final, min = 2,053,167
 N_final, max = 8,958,333

a) Fastest growth:
 n = log2(8,958,333 / 119,600) ≈ log2(74.9) ≈ 6.23
 g = 84 / 6.23 ≈ 13.5 hours

b) Slowest growth:
 n = log2(2,053,167 / 421,333) ≈ log2(4.87) ≈ 2.28
 g = 84 / 2.28 ≈ 36.8 hours

c) Mean-based estimate:
 Mean inoculum = (119,600 + 421,333) / 2 = 270,467
 Mean final = (8,958,333 + 2,053,167 + 3,962,167) / 3 = 4,990,222
 n = log2(4,990,222 / 270,467) ≈ log2(18.45) ≈ 4.21
 g = 84 / 4.21 ≈ 20.0 hours

##### +600 mV Condition

Final values:
 N_final, min = 360,583
 N_final, max = 523,483

a) Best-case scenario (minimum doubling time):
 n = log2(523,483 / 119,600) ≈ log2(4.38) ≈ 2.13
 g = 84 / 2.13 ≈ 39.4 hours

b) Worst-case scenario (no growth):
 n = log2(360,583 / 421,333) ≈ log2(0.856) ≈ -0.23 → no net growth

c) Mean-based estimate:
 Mean final = (360,583 + 523,483 + 458,983) / 3 = 447,683
 Mean inoculum = 270,467
 n = log2(447,683 / 270,467) ≈ log2(1.655) ≈ 0.731
 g = 84 / 0.731 ≈ 115.0 hours

##### Summary

| Electrode potential | Doubling Time Range | Mean-based Doubling Time |
| --- | --- | --- |
| +250 mV | 13.5 – 36.8 hours | ~20.0 hours |
| +600 mV | ≥39.4 hours (or no growth) | ~115.0 hours |

These calculations show that poised electrodes at +250 mV can support *Electronema aureum* GS growth rates comparable to those previously estimated for oxygen-respiring cable bacteria, while growth at +600 mV is substantially slower or negligible.

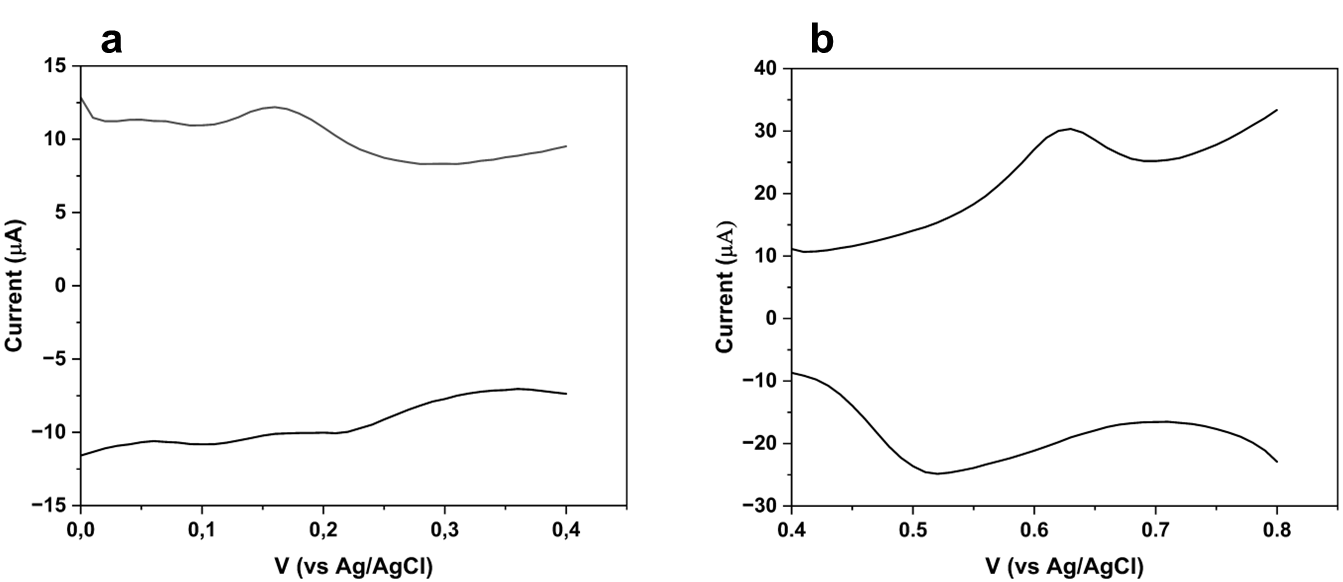

Figure S1. Scans across narrow potential windows revealed reversible peaks for *E.aureum* GS corresponding to the +250mV and +600mV peaks.

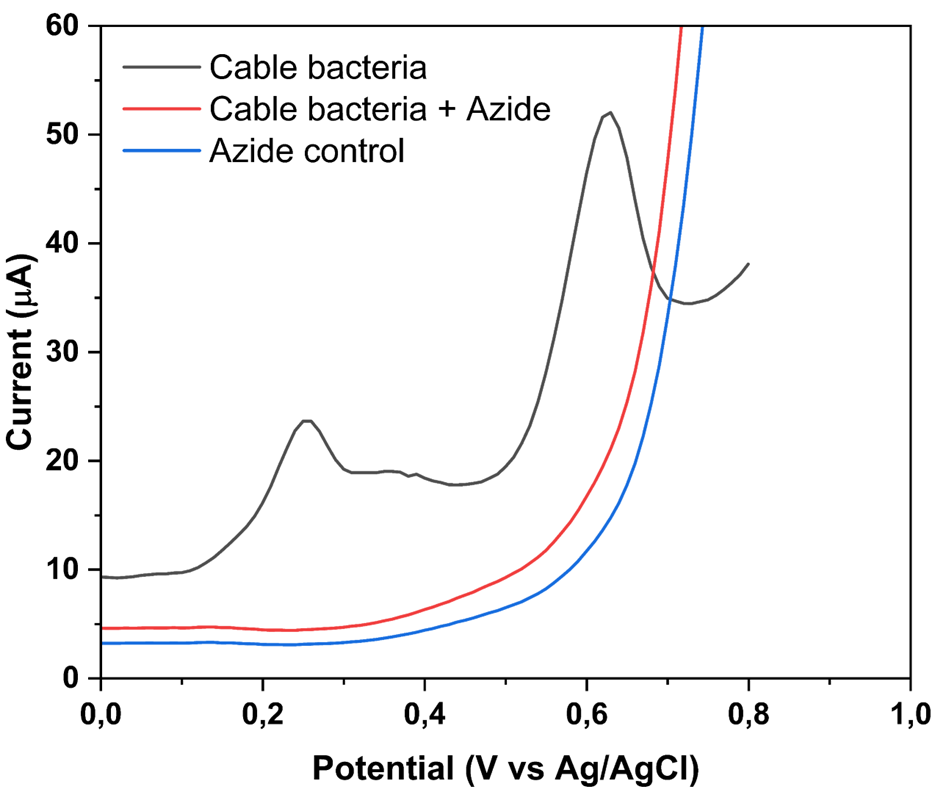

Figure S2. Effect of sodium azide on redox activity in *Electronema aureum* GS.
DPV profiles of cable bacteria before (black) and after (red) addition of 10 mM sodium azide. The redox peaks at ~+250 mV and ~+600 mV (vs Ag/AgCl) are abolished following azide treatment, indicating inhibition of redox-active moieties. Control measurements containing azide but no bacteria (blue) showed no redox features. These findings support the involvement of azide-sensitive, likely heme-containing outer-membrane proteins in extracellular electron transfer.

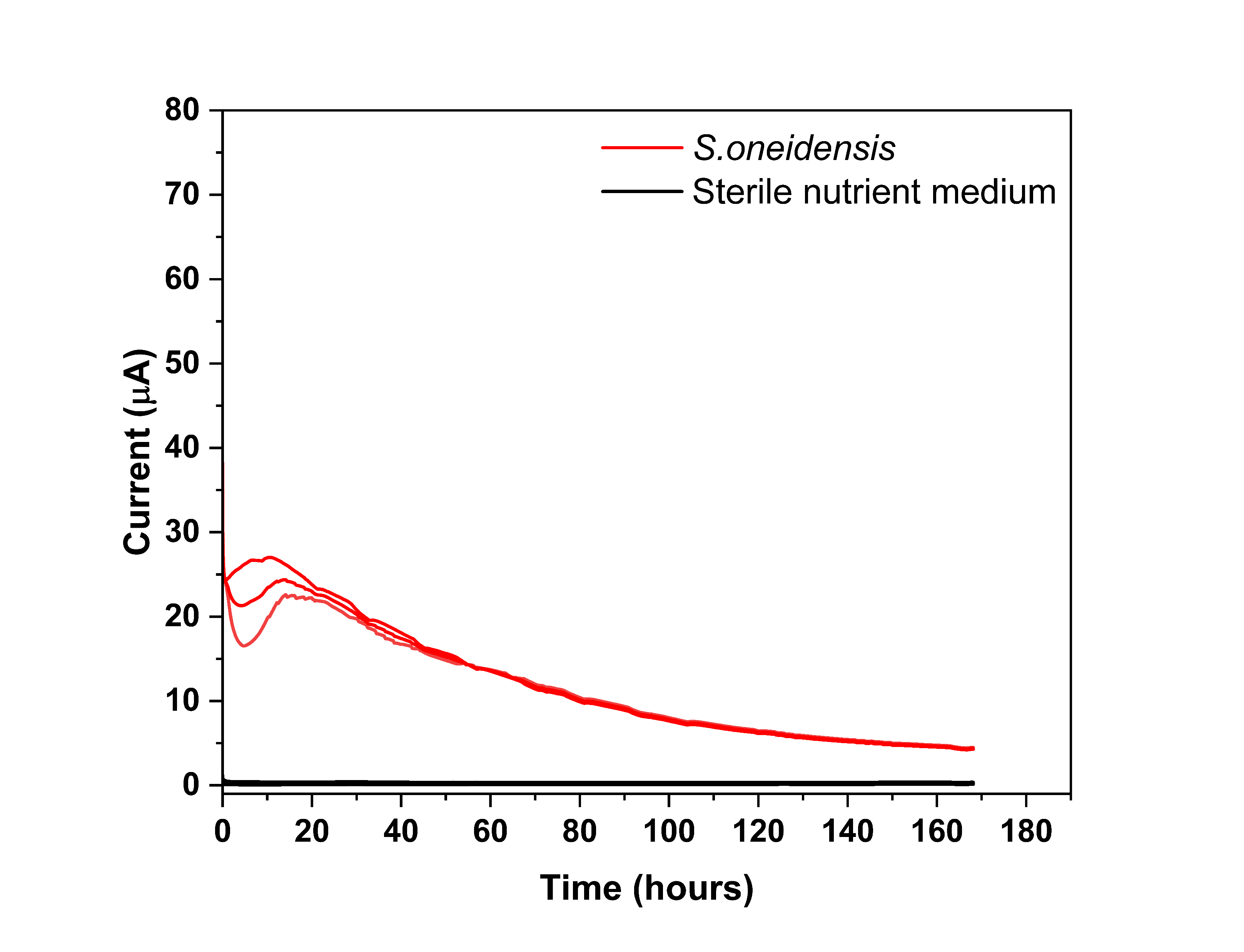

Figure S3. *Shewanella oneidensis* MR-1 poised at +600mV (vs Ag/AgCl) in bioelectrochemical systems did not generate increasing current over time.

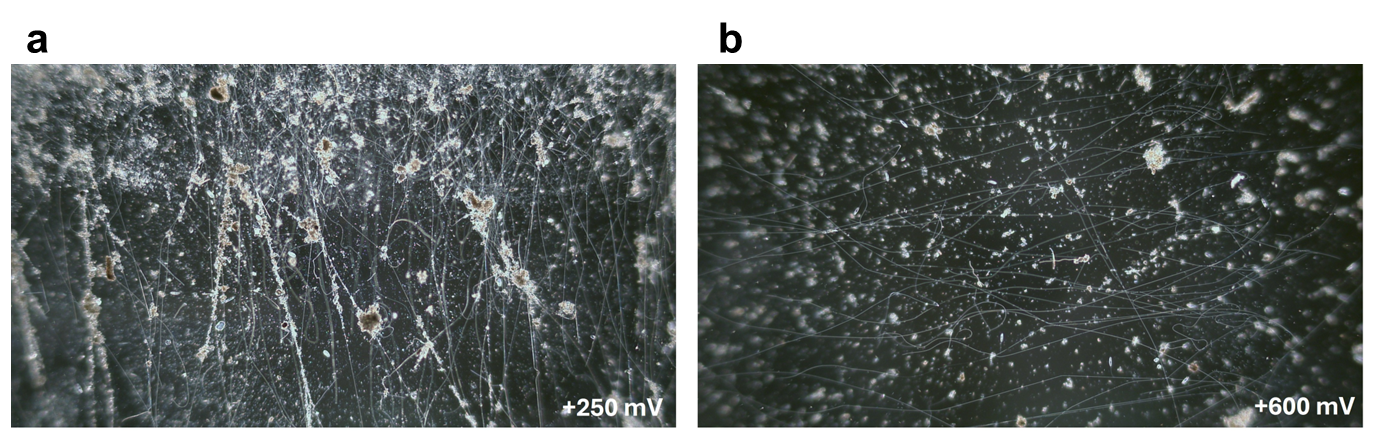

Figure S4. Living *E. aureum* GS filaments from bioelectrochemical systems poised at +250mV (a) and +600mV (b) respectively.

A

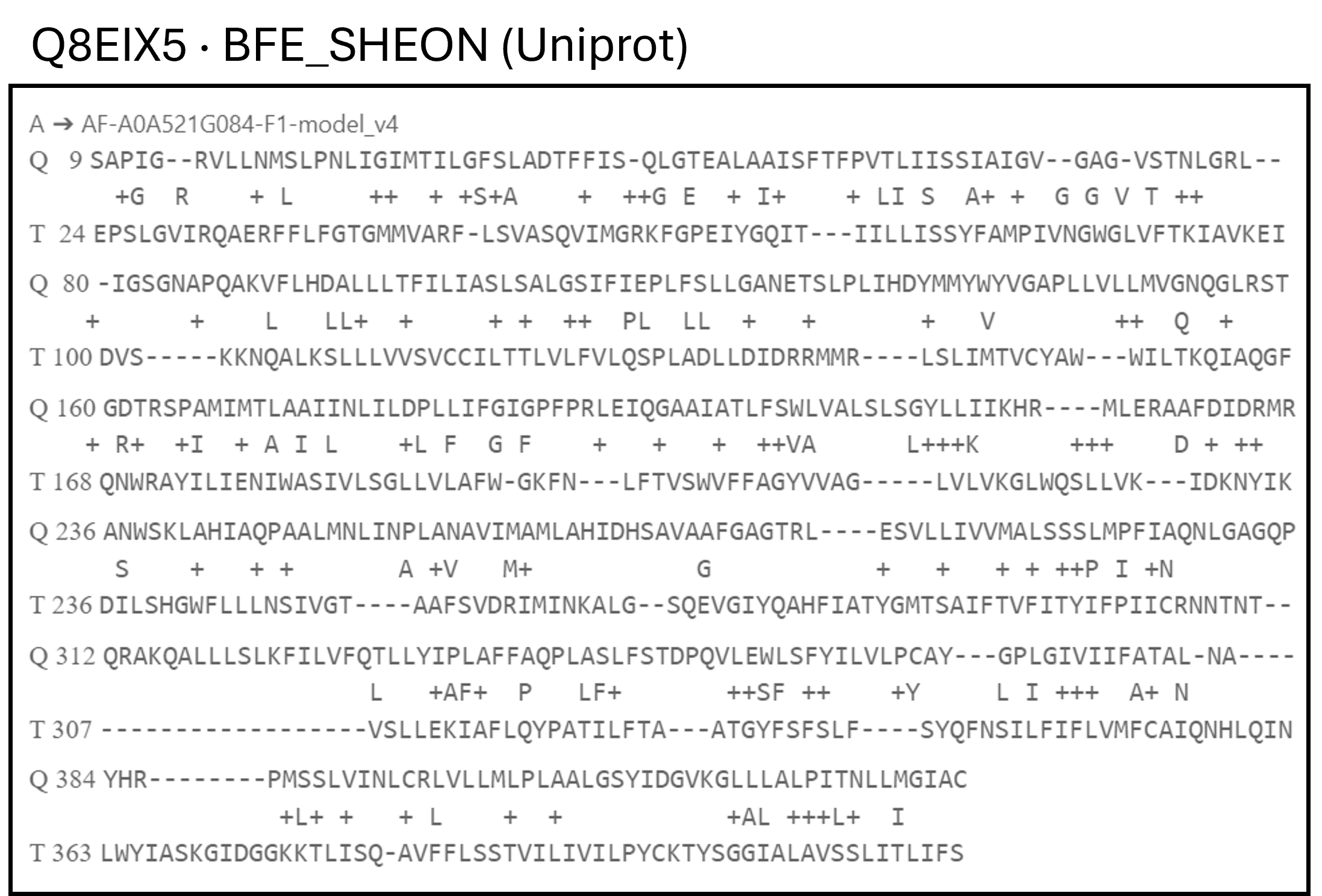

B

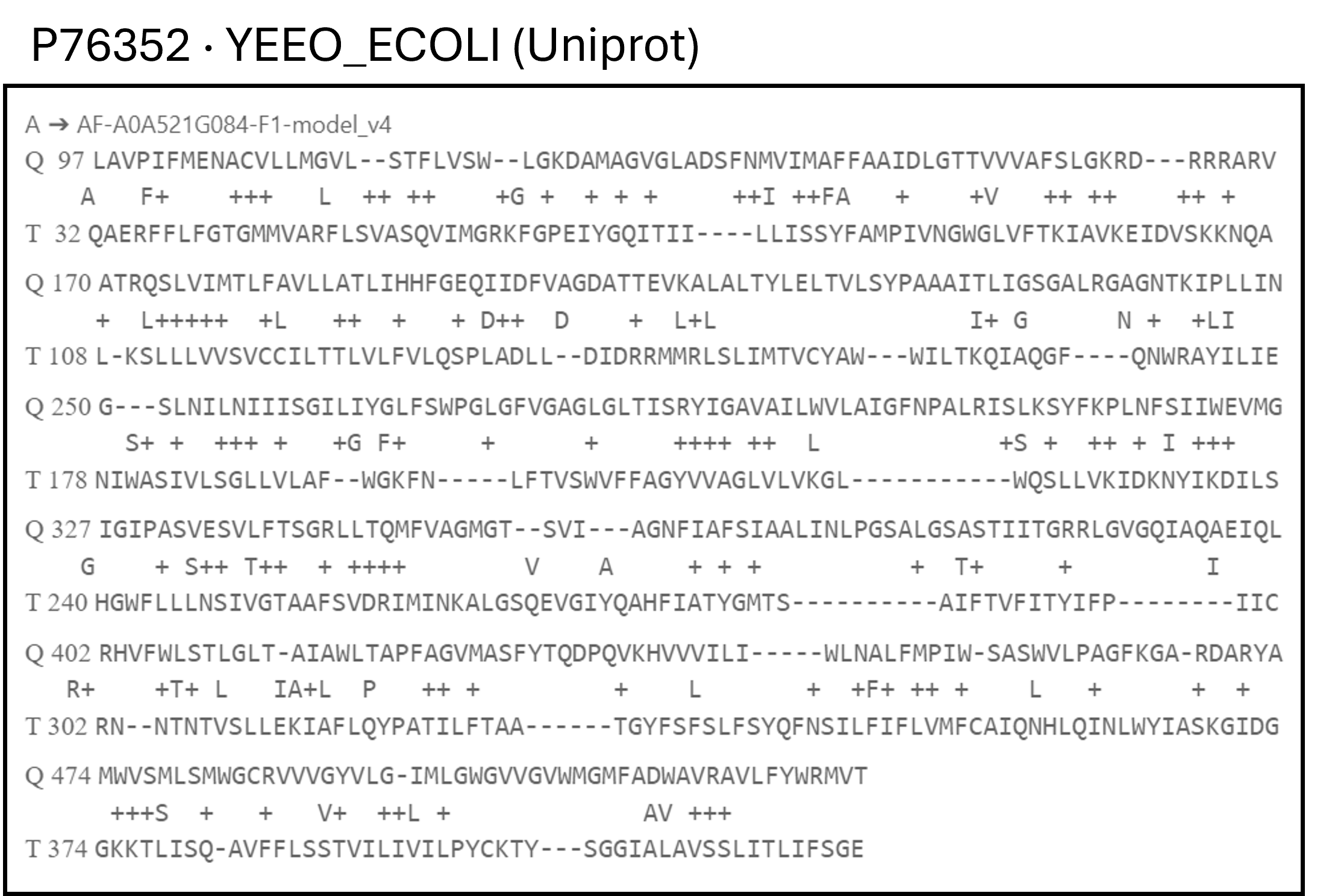

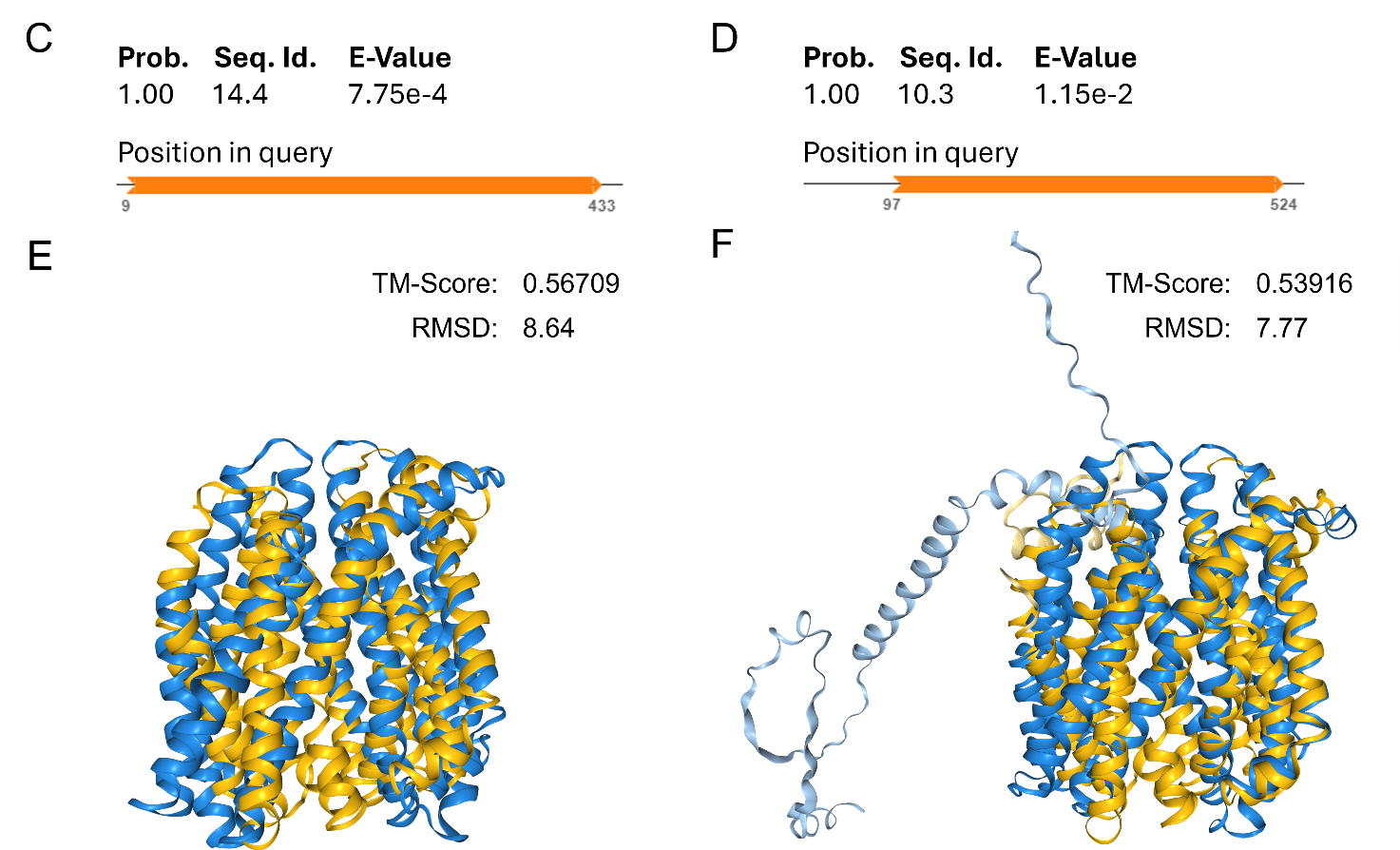

Figure S5. Alignments of sequence and structural homology for GS_00133 protein from Electronema aureum GS and FMN/FAD exporters. Overall sequence homology of the target GS_00133 to Bfe of and YeeO flavin exporters (queries) from Shewanella oneidensis strain MR-1 and E.coli strain K12 is low (A, B respectively). Structural homology search using AlphaFold protein models of Bfe and YeeO against Electronema aureum GS proteome (C and E, D and F respectively) matched both exporters exclusively to GS_00133. Alignments and homology searches were performed with Foldseek Search server. GS_00133; Uniprot accession number A0A521G084.

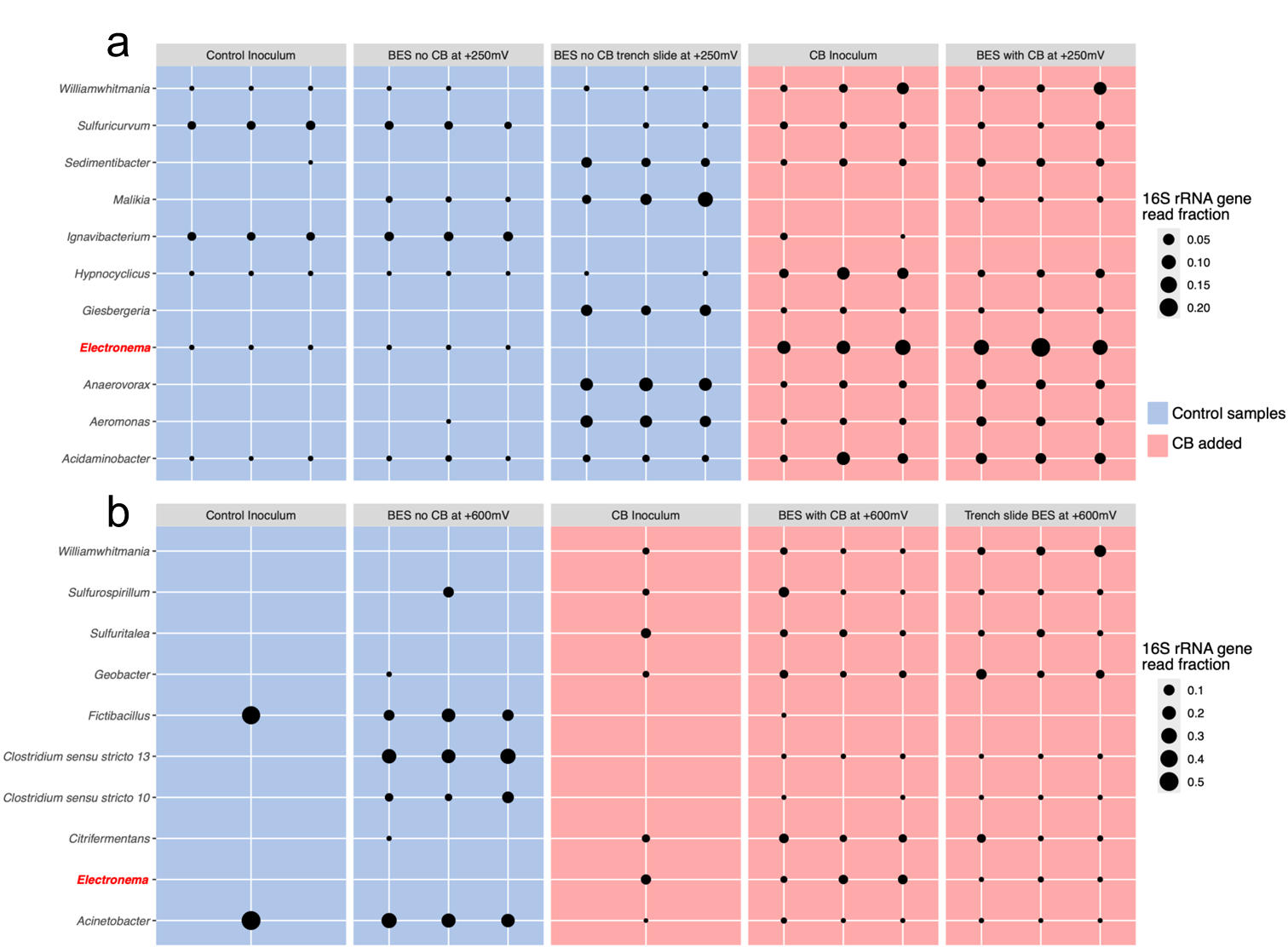

Figure S6. Microbial community composition in sediment BES. a. 16S rRNA gene sequencing of BES poised at +250 mV reveals enrichment of *E. aureum* GS in cable bacteria-inoculated sediments compared to controls. b. Similar analysis of BES poised at +600 mV shows survival of *E. aureum* GS.
